# Sterol regulatory element binding protein and Sp1 is required for transcriptional activation of a freshwater carnivorous fish (*Channa striata*) *elovl5* elongase

**DOI:** 10.1101/751032

**Authors:** Pei-Tian Goh, Meng-Kiat Kuah, Yen-Shan Chew, Hui-Ying Teh, Alexander Chong Shu-Chien

## Abstract

Long-chain polyunsaturated fatty acid (LC-PUFA) biosynthesis involves the activities of two groups of enzymes, the fatty acyl desaturases (Fads) and elongases of very long-chain fatty acid (Elovl). Fish are major source of beneficial n-3 LC-PUFA in human diet and there is a considerable interest to elucidate the mechanism and regulatory aspects of LC-PUFA biosynthesis in farmed fish species. The promoter of *elovl5* elongase, which catalyze the rate limiting reaction of adding two carbons to the C18 PUFA have been previously described and characterized from several marine and diadromous teleost species. We report here the cloning and characterization of the *elovl5* promoter from two freshwater fish species, the carnivorous snakehead fish (*Channa striata*) and zebrafish. Results show the importance of sterol regulatory element binding protein (Srebp) and the corresponding sterol responsive element (SRE) in the core regulatory region of both promoters. Mutagenesis luciferase and electrophoretic mobility shift assays confirm that SRE is indispensable for basal transcriptional activation in both species. In addition, several Sp1 binding sites located in close proximity with SRE were present in the snakehead promoter, with one having a potential synergy with SRE in regulating *elovl5* expression. The core *elovl5* promoter fragments of both species also directed *in vivo* expression in the yolk syncytial layer of developing zebrafish embryos. This study is the first functional promoter analysis of Elovl5 in freshwater teleost.

## Introduction

Long-chain polyunsaturated fatty acids (LC-PUFA) such as eicosapentaenoic acid (EPA, 20:5n-3), docosahexaenoic acid (DHA, 22:6n-3) and arachidonic acid (ARA, 20:4n-6), are fundamental for cellular membrane structure and function, eicosanoids synthesis, gene regulation and cellular signalling. In eukaryotes, LC-PUFA is generated from shorter chain polyunsaturated fatty acids (PUFA) through sequential actions of the fatty acyl desaturases (Fads) and elongases of very long-chain fatty acid (Elovl) enzymes. Elovl catalyse the rate-limiting reaction of elongating PUFA carbon chain with two carbon units (Nugteren, 1965, Leonard et al., 2004). In vertebrates, seven Elovl families (ELOVL1–7) have been described, with each family having specific preferences of fatty acid substrate (Leonard et al., 2004). ELOVL1, ELOVL3, ELOVL6 were shown to mainly elongate saturated fatty acids (SFA) and monounsaturated fatty acids (MUFA) while ELOVL4 is involved in elongation of very long-chain polyunsaturated fatty acids (Tvrdik et al., 2000, Naganuma et al., 2011, Moon et al., 2014). Both ELOVL5 and ELOVL2 are required the LC-PUFA biosynthesis pathway, where linolenic acid (LNA) and linoleic acid (LA) are converted to their respective LC-PUFA.

Adequate dietary intake of n-3 LC-PUFA is essential for optimal human health (Simopoulos, 1991). The capacity for *de novo* PUFA biosynthesis activities in many marine producers and subsequently, the conversion of these PUFA into LC-PUFA by consumers occupying different feeding niches, have led to marine products as the principal source of EPA and DHA for terrestrial inhabitants (Arts et al., 2001, Gladyshev et al., 2009). One conodrum in aquaculture is the need to reduce dependency on fish oil (FO) as a major aquafeed ingredient, despite its palatability and desirable LC-PUFA content (Tocher, 2015). The efficient use of the C-18 PUFA rich vegetable oils (VO) as an alternative for aquafeed requires an understanding of the species dietary LC-PUFA requirements. In addition, the LC-PUFA biosynthesis capacity of the animal, which inevitably depends on the repertoire of *fads* and *elovl* present in the genome, is an important deliberation. There is an increasing efforts to clone and characterise these enzymes from aquaculture candidate species (Leaver et al., 2008). Concomitantly, many of these studies also reported the upregulation of *fads* and *elovl* in response to limited availability of dietary LC-PUFA (Zheng et al., 2004, Carmona-Antoñanzas et al., 2014). Accordingly, efforts to characterise the upstream promoters in these genes began to transpire. So far, the promoter of *elovl5* have been isolated and characterised from salmon (Carmona-Antonanzas et al., 2016), orange grouper (Li et al., 2016), rabbit fish (Li et al., 2018) and golden pompano (Zhu et al., 2018). In contrast, there is no investigation on any freshwater species. Given the difference in the LC-PUFA biosynthesis capacity between several marine and freshwater teleost, it is important to decipher the regulatory elements driving transcription in promoters of Elovl from freshwater aquaculture candidate species.

The striped snakehead (*Channa striata*; Bloch, 1793) is a freshwater carnivorous teleost fish with relatively high tissue deposition of DHA and is farmed throughout Southeast Asia (Wee, 1982). We have previously showed that *C. striata* possess a complete set of Fads and Elovl for production of LC-PUFA from C18-PUFA, inclusive of an *elovl5* capable of C18-C20 PUFA elongation (Kuah et al., 2015, Kuah et al., 2016). The zebrafish is a useful model to understand the developmental role of lipid and fatty acids (Holtta-Vuori et al., 2010, Quinlivan and Farber, 2017). Relevant to our work, several zebrafish *fads* and *elovl* ortholog have been identified and characterised in terms of function and spatio-temporal expression pattern (Hastings et al., 2001, Agaba et al., 2004, Tan et al., 2010, Monroig et al., 2010). We describe here the cloning and characterisation of the snakehead and zebrafish *elovl5* promoters. We demarcated the regulatory elements for driving transcription and also traced the *in vivo* expression of both promoter sequences in developing zebrafish embryos.

## Materials and methods

### Fish maintenance and breeding

Juvenile snakehead fish were raised in outdoor cement tanks (inner dimension: 130×98×280 cm) at temperature of 29 ± 1 °C, under natural photoperiod and fed ad libitum with commercial pellets (Star Feedmills, Malaysia). Wild type zebrafish (AB line) were maintained in the ZebTEC Stand Alone System (Tecniplast, USA). Fishes were kept on a 13:11 h light/dark cycle at 28.5 °C and fed twice daily with commercial micro pellet (Aquadene, Malaysia), frozen bloodworms and *Artemia* nauplii. Breeding and embryo collection were carried as previously (Tay et al., 2018). Handling and sacrifice of *C. striata* and *D. rerio* were in accordance to the guidelines by the Animal Ethics Committee, Universiti Sains Malaysia (PA/ACSC/002/2011) with accompanying approval

### Cell line maintenance and subculture

Zebrafish liver cell line, ZFL (ATCC® CRL-2643™) (American Type Culture Collection, USA) was maintained in complete growth medium at 28 °C following the ATCC protocol. Routine subculture was performed upon 80 to 90% confluence in a 25 cm^2^ flask.

### Cloning of the snakehead and zebrafish elovl5 promoters

DNA extraction from fish liver tissue was conducted using the QIAmp^®^DNA Mini Kit (Qiagen, Germany). For snakehead fish *elovl5* promoter, the GenomeWalker^®^ Universal kit was used. All primers are listed in Supplementary 1. PCR amplification of promoter fragments were conducted using the Advantage 2^®^ PCR enzyme system (Clontech, USA) according manufacturer’s protocol. For fragments that are shorter than 1 kb, the thermal cycling program comprised an initial denaturation step for 1 min at 95°C, followed by 35 amplification cycles for 30 s at 95°C and 1 min at 68°C, and a final extension step for 1 min at 68°C. For longer fragments, the annealing and final extension steps was extended to 3 min at 68°C, respectively. The PCR products were cloned and ligated into a restricted pGL3-Basic luciferase reporter vector (Promega, USA). For 5’-deletion analysis of snakehead *elovl5* gene promoter, six different length luciferase promoter-reporter plasmids were constructed using primers containing a restriction site for *Sac* I and *Bgl* II, respectively.

For zebrafish, a DNA fragment of 2794 bp, corresponding to the −2592/+202 bp region of the transcription start site of zebrafish *elovl5* was isolated utilising the *i*-Taq Plus DNA Polymerase kit as per manufacturer’s instructions (iNtRON, Korea). DNA was extracted from adult zebrafish using CTAB lysis buffer, and subjected to a PCR mixture as the template, coupling with a forward primer containing the restriction site for *Kpn*I, and a reverse primer containing the restriction site for *Xho*I (Supplementary 1). The amplified PCR products were analysed on 0.75 % (w/v) agarose gel, stained with SYBR Safe DNA Gel Stain and visualized under UV transilluminator. The PCR products were cloned into the pGL3-Basic luciferase reporter vector as described above. Four promoter fragments were constructed for 5’-deletions analysis (Supplementary 1).

### Transient transfection and dual luciferase reporter assay

Transfection assay to assess the capacity of different promoter fragments to drive transcription activities was carried out using the Lipofectamine^®^ transfection reagent (Invitrogen). Approximately 2 × 10^5^ ZFL cells were seeded into 96-well plate 24 hr before transfection. Next, 0.25 µl of lipofectamine was diluted in 6 µl of serum-free medium (Opti-MEM^®^ I), with the mixture incubated for 5 minutes at room temperature. These cells were transfected with 150 ng of the pGL3 promoter-luciferase reporter plasmid of interest, 50 ng of *Renilla* pRL-SV40 internal control plasmid and 50 ng pcDNA3.1-*Danio rerio* Srebp. Subsequently, 50 µl of serum-free medium was added. The mixture was overlaid into the 96-wells plates and incubated at 28.5°C for 5 hours, followed by replacement with 125 µl of fresh complete DMEM and further incubated for 24 hours. Luciferase assay was carried out on transfected cells using the Dual-Glo^®^ Luciferase Assay System (Promega, USA). Prior to the 5’-deletion assay, the effect of zebrafish Srebp1 and Srebp2 on luciferase activities was compared by co-transfecting ZFL cells with promoter reporter plasmids and expression plasmid containing either Srebp.

### Site-directed mutagenesis

Site-directed mutagenesis was carried out on putative binding elements in snakehead and zebrafish promoters using the Mutadirect^®^ Site Directed Mutagenesis Kit (Intron, Korea) Two complementary oligonucleotides containing the desired mutation sequence were designed were designed using PrimerX (http://www.bioinformatics.org/primerx/) and shown in Supplementary 1. For each reaction, 5 μl of Muta-DirectTM Reaction Buffer, 2 μl of dNTP mixture, 1 μl of each forward and reverse mutagenic primers (10 pmol/μl), 2 μl of plasmid DNA (10 ng/μl), 1 μl of Muta-DirectTM Enzyme (2.5 U/ μl) and 38 μl of sterile water were subjected to PCR amplification programmed with an initial denaturation cycle for 30 s at 95°C, followed by 18 cycles of denaturation for 30 s at 95°C, annealing for 1 min at 55°C, and extension for 1 min/kb of plasmid length. Subsequently, the non-mutated parental plasmid DNA templates were digested with MutazymeTM Enzyme for 90 min at 37°C. Newly transformed bacterial colonies were selected for overnight propagation in LB medium. The mutated plasmids were then purified from the bacterial cells and verified through sequencing. Luciferase assay for the mutated fragments are as above.

### Electrophoretic mobility shift assay (EMSA)

EMSA was carried out using LightShift^®^ Chemiluminescent EMSA Kit (Thermo Scientific, USA). Labelling of the 3′-end of each oligonucleotide complementary strand with biotin was conducted with the Biotin 3′ End DNA Labelling Kit (Thermo Scientific, USA). Labelled double-stranded oligonucleotides and non-labelled competition double-stranded oligonucleotides were generated at least 1 h prior to EMSA. Nuclear extract was prepared from transfected ZFL cells using the NE-PER^®^ Nuclear and Cytoplasmic Extraction Reagents (Thermo Scientific, USA). For the negative control binding reaction, nuclear extract was excluded, while for competition binding reactions, 100-fold molar of non-labelled oligonucleotides was included. The DNA-protein binding complexes were separated on a 5% (v/v) non-denaturing polyacrylamide gel at 4 °C (Bio-Rad Mini-PROTEAN, USA). Chemiluminescent Nucleic Acid Detection Module (Thermo Scientific, USA) was used for detection.

### In vivo observation of snakehead and zebrafish *elovl5* promoters in zebrafish embryos

For visualization the *in vivo* expression of both respective *elovl5* promoters, promoter-pZsGreen1-1 GFP reporter plasmids were constructed through amplification of desirable promoter regions using forward and reverse primers containing *Xho*I and *BamH*I restriction sites (Supplementary 1). The reporter plasmids were linearised with the restriction enzymes *XhoI* and *NotI* prior to microinjection. Concentration of reporter plasmid was adjusted to 10 ng/μl with dH2O containing 1X Danieau’s buffer and 0.2% phenol red and approximately 4.6 nl of solution was delivered into one-cell stage embryos using the Nanoliter 2000 Microinjector (World Precision Instrument, USA). Injected embryos were incubated in E3 medium at 28.5 °C and periodically monitored under the MVX10 Fluorescence Macro Zoom Microscope equipped with ColorView III Soft Imaging System (Olympus, Japan) until 120 hpf.

## Results

### Snakehead and zebrafish *elovl5* promoters

A snakehead *elovl5* DNA fragment containing a 5’-UTR and an upstream promoter region of 5905 bp was obtained by combining three DNA fragments (Supplementary 2). The 5’-UTR region consists of four exons and three introns. The first three exons, 1A, 1B and 1C are non-coding exons. Three putative transcription start sites (TSS) were identified. Subsequently, the region adjacent to exon 1A (−2497/+46 bp) was cloned for further characterization. A 2.8 kb 5’ flanking sequence of the zebrafish *elovl5* promoter region was isolated (Supplementary 3). A putative transcription start site was located at exon 1. Exon 2 is separated from the first exon by approximately 4.2 kb of intron.

5’ deletion and mutagenesis luciferase assays of the snakehead and zebrafish *elovl5* promoters.

Co-transfection of either the snakehead 2.5kb *elovl5* or zebrafish 2.8kb *elovl5* promoter-luciferase reporter plasmid with an expression plasmid containing the nuclear form of zebrafish Srebp1 resulted in significantly higher reporter activities than cells transfected with empty pcDNA3.1 expression plasmid or the zebrafish Srebp2 protein (Fig. 1). Therefore, ensuing luciferase reporter transactivation assays were carried out with overexpression of zebrafish nSrebp1 protein.

**Figure 1.**
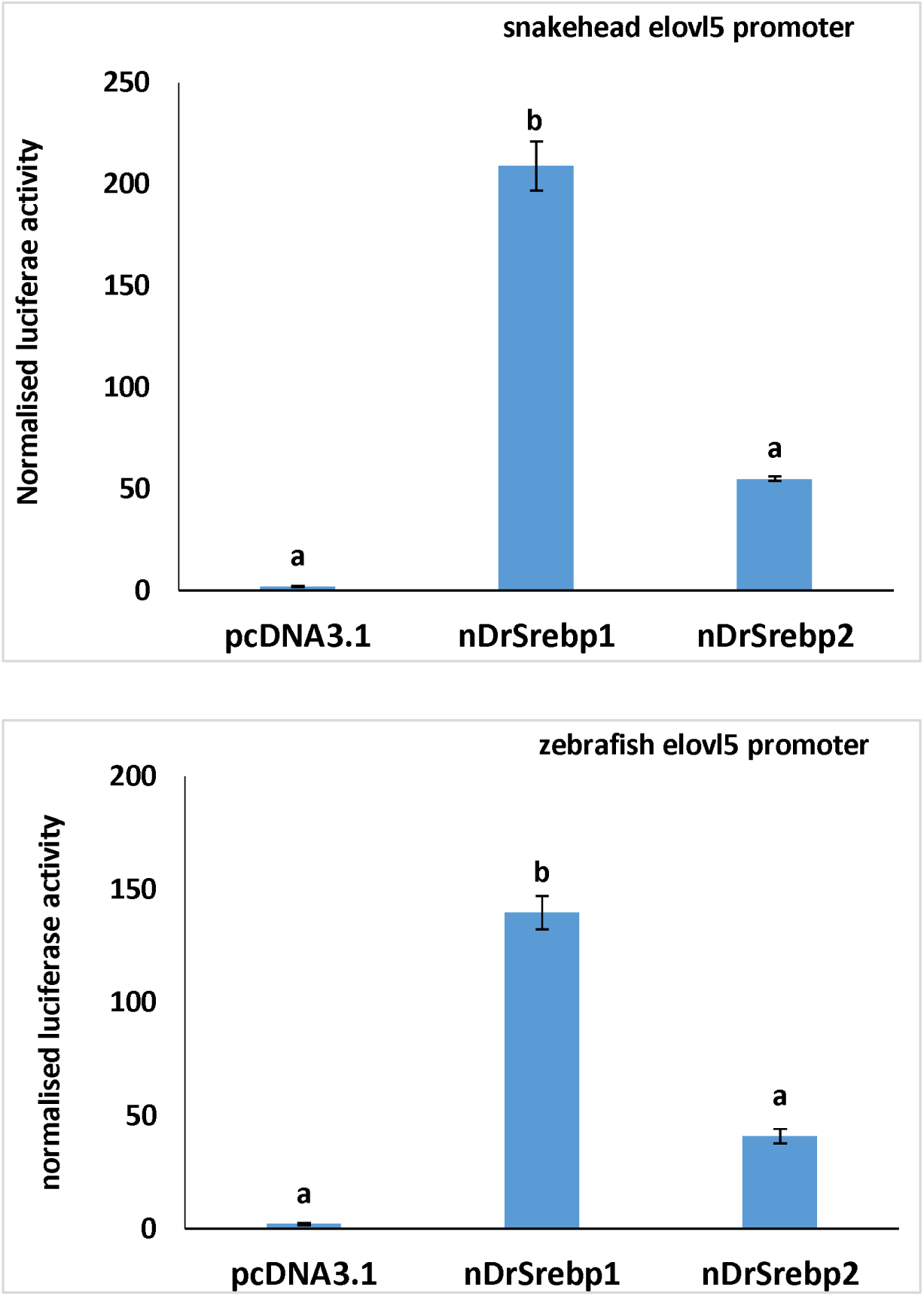
Luciferase activities of snakehead 2.5kb *elovl5* and zebrafish 2.8kb *elovl5* promoters when nuclear form of zebrafish Srebp 1 or Srebp 2 proteins are over-expressed in ZFL cells. Co-transfection of cells were carried out with pGL3 promoter-luciferase reporter plasmid fused with the snakehead *elovl5* (−2497/+46 bp) or zebrafish *elovl5* promoter (−2592/+202 bp) and pcDNA3.1 expression plasmids fused with either zebrafish Srebp1 or Srebp2 as described previously. Results represent normalised luciferase activity of *elov5* promoter in the ZFL cells in relative to empty pGL3-basic plasmid.

For the snakehead *elovl5* promoter, 5’-deletion analysis revealed that maximal promoter activity was observed with fragment consisting 1460 nucleotides upstream of the TSS (Fig. 2). Deletion of region between −1460 to −474 bp significantly reduced the promotewr transcriptional activity (*P* < 0.05). Additional deletions upstream of this fragment did not further reduce transcriptional activities. Collectively, these results indicated that the snakehead *elolv5* promoter region between the 161/+46 bp may contain core regulatory elements. *In silico* analysis of this region with P-Match (www.gene-regulation.com) reveals three putative binding sites for the ubiquitous transcription factor Sp1 and a sterol regulatory element (SRE), binding site (Supplementary 4). Mutating the SRE completely abolished transcription activities of the promoter (Fig. 3A). Sequence mutation of one of the Sp1 site significantly lowered the luciferase intensity as compared to the control fragment. Interestingly, obliterating the +11 to +21 Sp1 site adjacent to the SRE increased luciferase activities.

**FIGURE 2:**
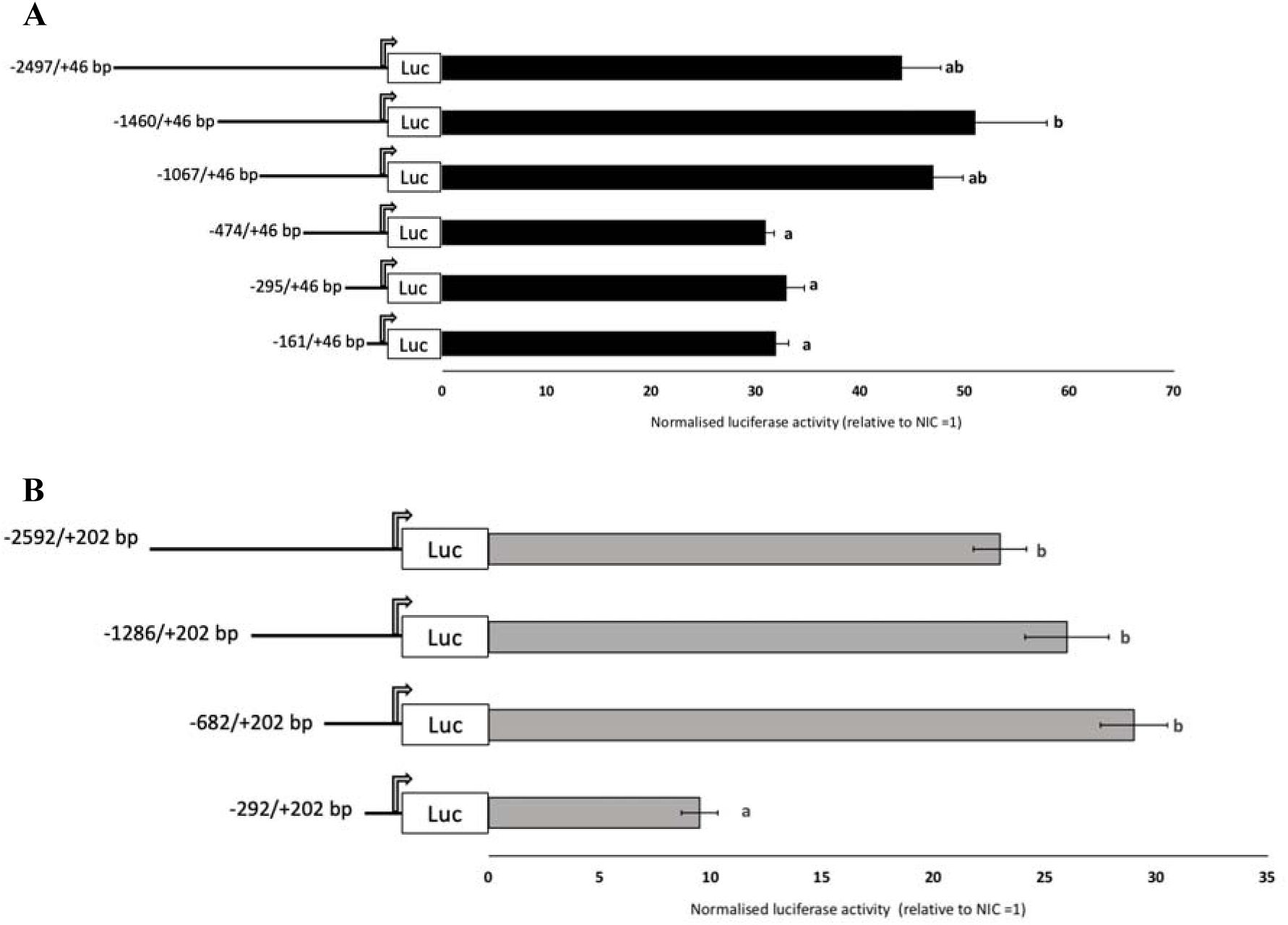
5’-deletion analysis of (A) 2.5kb snakehead *elovl5* promoter (B) −2.8kb zebrafish *elovl5* promoter. Constructs are named according to position from TSS (arrow). Dual-reporter luciferase assay was conducted using ZFL cells co-transfected with promoter-luciferase reporter plasmid, nSrebp1 expression plasmid and *Renilla* luciferase reference plasmid pRL*-*SV40. Luciferase activity (mean ± S.E) was normalized to empty pGL3-Basic plasmid. Mean values were subjected to one-way ANOVA and ranked with Tukey’s test (P<0.05).

**FIGURE 3.**
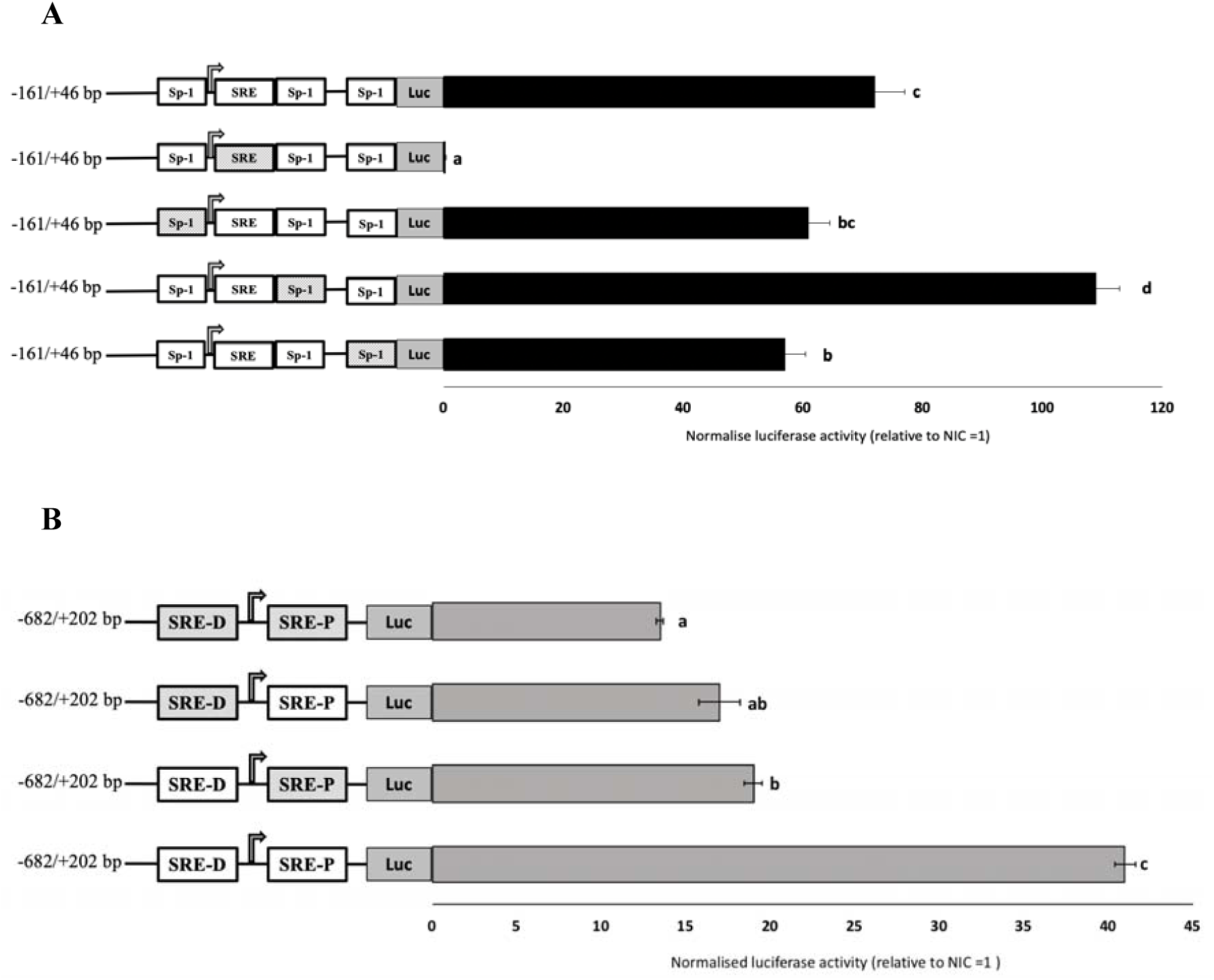
Mutagenesis analysis of (A) −161/+46 bp snakehead *elovl5* promoter (B) 682/+202 bp zebrafish *elovl5* promoter. Mutated putative binding site are shaded. TSS is represented by arrow. Dual-reporter luciferase assay was conducted using ZFL cells co-transfected with promoter-luciferase reporter plasmid, nSrebp1 expression plasmid and *Renilla* luciferase reference plasmid pRL*-*SV40. Luciferase activity (mean ± S.E) was normalized to empty pGL3-Basic plasmid. Mean values were subjected to one-way ANOVA and ranked with Tukey’s test (P<0.05).

For zebrafish, a significant drop in luciferase readout was obtained when the promoter was truncated to −292/+202 bp, which indicates the presence of necessary binding elements for basal transcription within the −682/+202 bp fragment (Fig. 2B). Bioinformatic analysis reveals two putative SRE sites located at −376 and +186bp (Supplementary 4). Mutating either of the SRE binding site or both sites resulted in significantly lower transcriptional activities than the intact fragment (Fig. 3B).

Binding of zfl nuclear proteins to sre sites of snakehead and zebrafish elovl5 promoters. EMSA showed the formation of four DNA-protein complexes when biotin-labelled oligonucleotides containing the putative snakehead SRE sequence 5’-GCTCAGACGGG-3’ was incubated with nuclear extract of ZFL cells (Fig. 4). The specificities of these bindings were validated when oligonucleotides with mutated SRE sequences or wildtype oligonucleotide incubated together with excessive non-labelled oligonucleotide did not yield any DNA-protein complex.

**FIGURE 4:**
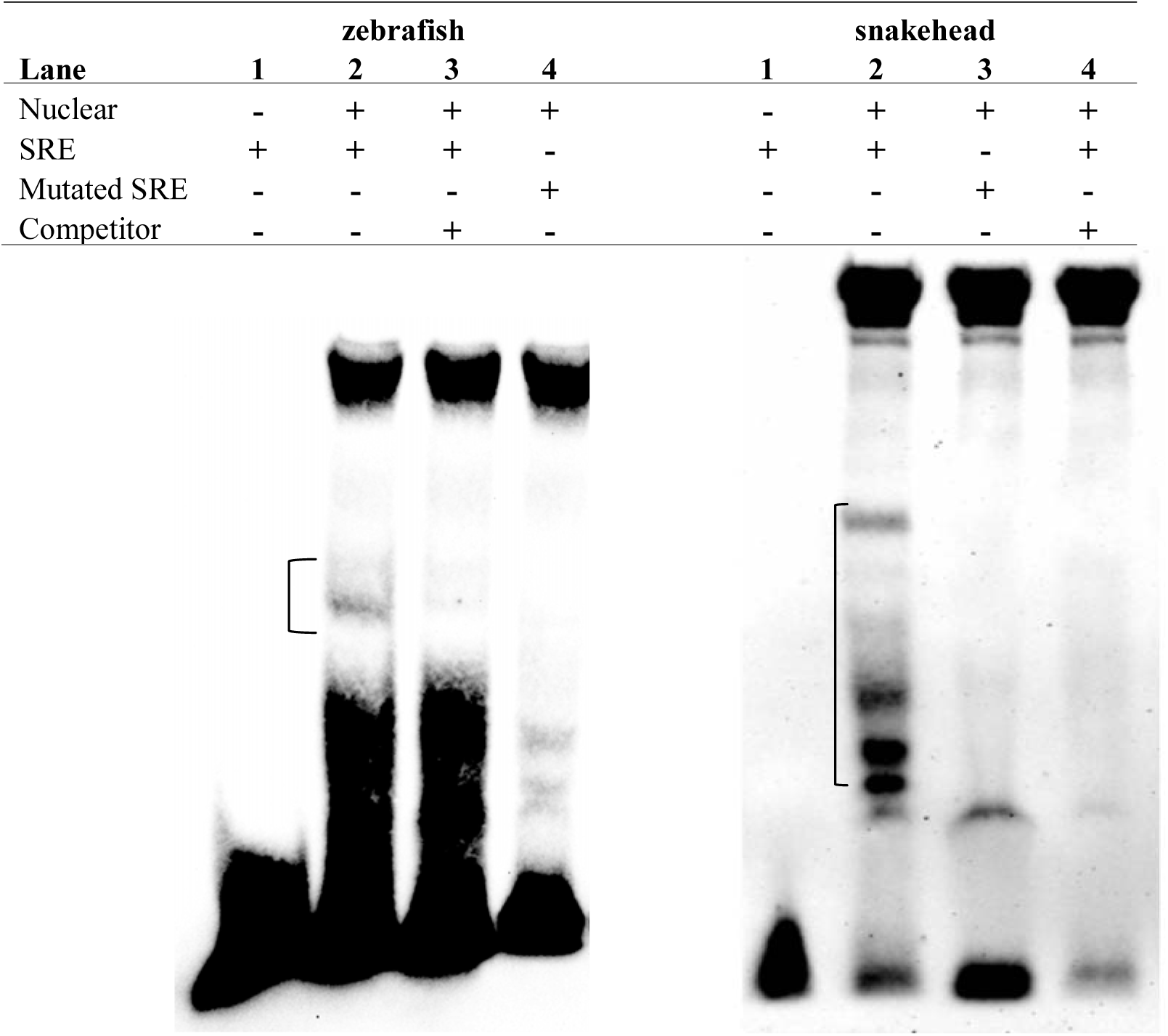
Binding of ZFL proteins to the potential zebrafish and snakehead fish *elovl5* SRE sites. EMSA was conducted with biotin-labelled oligonucleotides containing SRE of respective promoter species and nuclear extract of ZFL cells. Formation of DNA binding-protein complex (bracket) was verified with competitive assays using excess unlabeled oligonucleotides or mutated SRE oligonucleotides of respective promoter species.

As for zebrafish, DNA-protein complex was formed when ZFL cells nuclear extract was incubated with biotin-labelled oligonucleotide probe containing the proximal SRE sequence. There was no band when probe was incubated with excessive unlabelled proximal SRE oligonucleotide. Incubation of nuclear extract with labelled mutated promixal SRE probe resulted in non-specific binding.

### Transient expression of snakehead and zebrafish *elovl5* promoters in developing zebrafish embryos

Microinjection 682/+202 bp zebrafish *elovl5* promoter into developing embryos (n = 198) resulted in specific GFP signalling in YSL (10.6%) from 24 hpf until 96hpf before declining (Fig. 5A-C). Injection of the −295/+46 bp fragment of the snakehead *elovl5* fused to GFP reporter plasmid resulted in 30% of injected embryos (n= 133) expressing GFP signal in the yolk syncytial layer (YSL), at 48-96hpf stage, before gradually subsiding at later stages (Fig. 5D).

**FIGURE 5:**
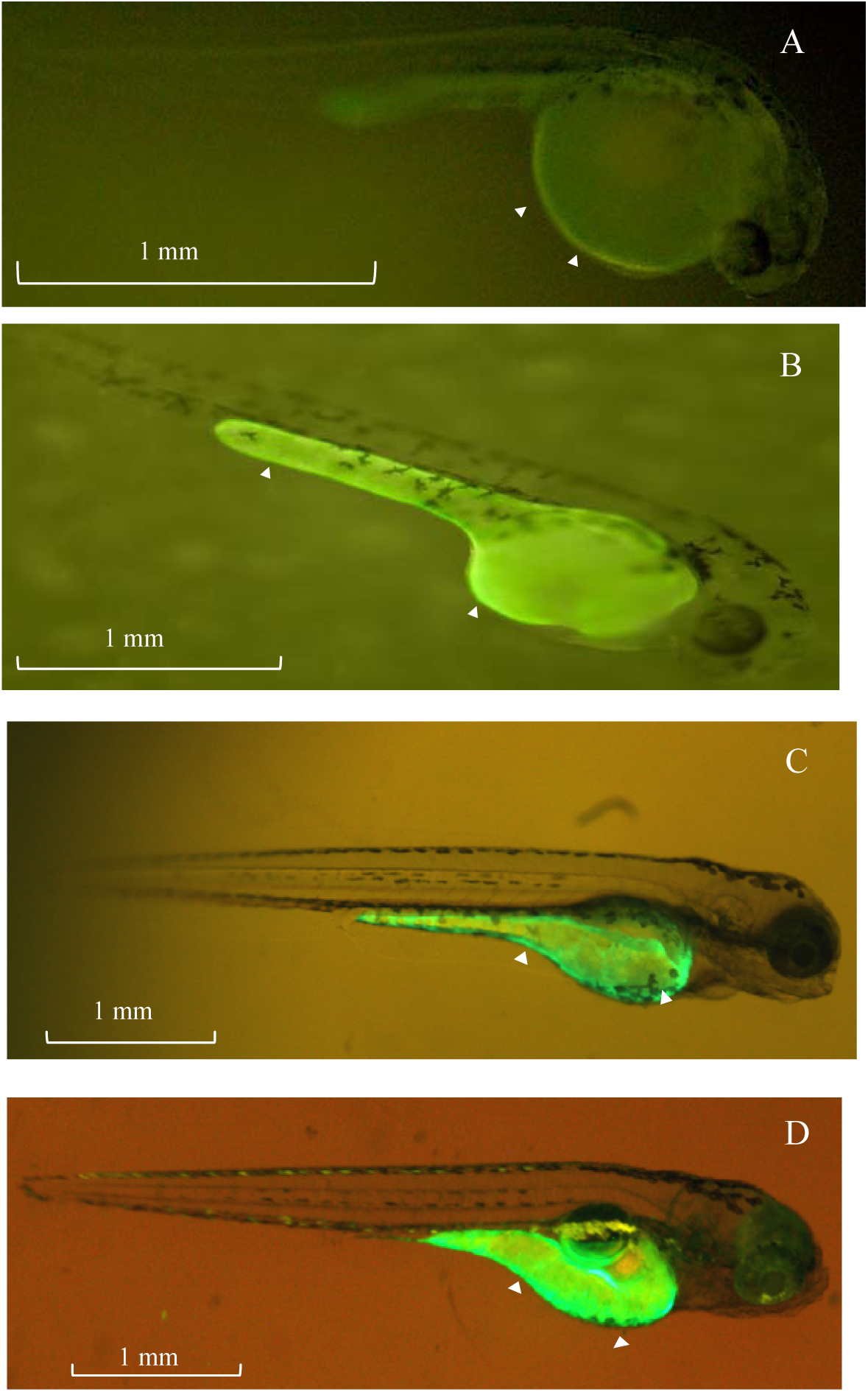
Transient GFP expression of *elovl5* promoter fragments (−682/+202 bp zebrafish; −295/+46 bp snakehead) in different stages of zebrafish embryos. (A) 24 hpf zebrafish *elovl5*, (B) 50hpf zebrafish *elovl5* (C) 72 hpf zebrafish *elovl5* (D) 72hpf snakehead *elovl5*. Arrowheads depict expression in yolk syncytial layer.

## Discussion

Orthologs of *elovl5* have been isolated from numerous marine, freshwater and diadromous teleost species (Agaba et al., 2004, Agaba et al., 2005, Leaver et al., 2008, Kuah et al., 2015). In tandem, diets with low levels of LC-PUFA are reported to upregulate the expression of *elovl5* in liver and intestinal tissues, to compensate for low LC-PUFA availability (Zheng et al., 2004, Zheng et al., 2005, Kuah et al., 2015). The participation of Fads and Elovl in LC-PUFA biosynthesis partly depends on the binding of specific transcription factors on their respective response elements in the promoter regions, and these activities are tightly regulated by hormones and nutritional status. To date, literature on teleost *elovl5* promoters are limited to marine and diadromous species (Li et al., 2016, Carmona-Antonanzas et al., 2016, Li et al., 2018, Zhu et al., 2018). The snakehead, a freshwater carnivorous fish, is an important candidate for intensive aquaculture in tropical countries while the zebrafish, a useful model for studying the function and regulation of Elovl and Fads (Hastings et al., 2001, Agaba et al., 2004, Tan et al., 2010, Monroig et al., 2010, Tay et al., 2018). Here, we studied the sequences of the putative *elovl5* promoters of snakehead and zebrafish, and characterised their ability to drive reporter expression *in vitro* and *in vivo*.

Results from the deletion and mutagenesis luciferase reporter assays delineate transcriptional activating roles for SRE in snakehead and zebrafish *elovl5* promoters. SREs are binding site for Srebp, which is a conspicuous regulator of cholesterol, fatty acid, triacylglycerol and phospholipid biosynthesis (Eberlé et al., 2004, Sampath and Ntambi, 2005). In mammals, three SREBP isoforms (SREBP-1a, SREBP-1c and SREBP-2) exist. Among them, SREBP-1a activates target genes involved in synthesis of triglycerides, cholesterol and fatty acids. SREBP-1c and SREBP-2 activities are more restricted, with the former mainly targeting genes involved in fatty acid synthesis including Elovl, Fads and fatty acid synthase (Horton et al., 2002, Qin et al., 2009). Teleost only possess a single form of Srebp1 and Srebp2, respectively. Our results showed that in ZFL cells, Srebp1 promotes higher transcriptional activities in snakehead and zebrafish *elovl5*, as compared to Srebp2. In Atlantic salmon, Srebp2 resulted in higher activities of *elovl5b* while both Srebp 1 and Srebp 2 equally activate *elovl5a* expression (Carmona-Antoñanzas et al., 2014, Carmona-Antonanzas et al., 2016). Recently, knockout of *elov2* in Atlantic salmon suggest a role of Srebp1 in feedback regulation of LC-PUFA biosynthesis (Datsomor et al., 2019). Atlantic salmon *fads2* promoter also contain a highly conserved SRE site which is regulated by both Srebp1 and Srebp2 (Zheng et al., 2009, Carmona-Antoñanzas et al., 2014). On the other hand, the transcriptional regulation of zebrafish *fads2* seemed to depend more on Srebp2 (Tay et al., 2018).

Both snakehead and zebrafish *elovl5* promoters contain SRE sites, which when obliterated, resulted in lower transcriptional activities. Additionally, overexpression of Srebp1 also induced a significant increase in promoter-driven luciferase activities of both species. This strongly suggest the involvement of Srebps in regulation of freshwater teleost *elovl5*, although the exact mechanism remains to be clarified. In liver, transcription mediators such as Liver X Receptor (LXR) or SREBP utilizes the intracellular levels of PUFAs to regulate PUFA homeostasis, including biosynthesis activities. Both EPA and DHA has been proven to inhibit the regulated intramembrane proteolysis (RIP) of the nascent SREBP-1c, resulting in an impeded level of transcriptionally active nuclear fragment (Takeuchi et al., 2010, Deng et al., 2015). Srebp-1 was also proposed to be the mediator for LC-PUFA biosynthesis in Atlantic salmon via activation in limited LC-PUFA availability condition or inhibition when fish are fed with n-3 LC-PUFA rich diet (Datsomor et al., 2019). In both snakehead and zebrafish, treatment with diets containing limited amounts of DHA or EPA induced the expression of hepatic *elovl5* (Kuah et al., 2015, Jaya-Ram et al., 2008, Jaya-Ram et al., 2016). In mouse, the suppression of Elovl5 by PUFA is mediated by SREBP-1c (Qin et al., 2009). Another postulated pathway involves the LXR, where stimulation of LXR by ligands stimulates the production of SREBPs (Qin et al., 2009, Carmona-Antoñanzas et al., 2014). LC-PUFA are also capable of inhibiting the binding of LXR to the SREBP-1c promoter, downregulating the transcriptional activities (Yoshikawa et al., 2002). In two marine fish species, *Larimichthys crocea* and *Epinephelus* coioides, and Atlantic salmon, LXR and SREBP-1 are involved in regulation of *elovl5* for LC-PUFA biosynthesis (Minghetti et al., 2011, Carmona-Antoñanzas et al., 2014, Li et al., 2016, Li et al., 2017). The hepatocyte nuclear factor 4 (Hnf4α) another lipogenesis regulator, is recently shown to be an essential regulator of LC-PUFA biosynthesis in a marine herbivorous fish *Siganus caniliculatus* by positively regulating both Fads and Elovl (Li et al., 2018, Wang et al., 2018, Dong et al., 2018). Furthermore, post-transcriptional regulation of *fads* and *elovl5* involving the enhancement of Srebp1 through microRNAs and an insulin-induced gene was also shown in the same species (Sun et al., 2019).

Besides neofunctionalization of LC-PUFA biosynthesis genes, having different regulatory pathways presumably increase the ability to adapt to environment with limited availability of LC-PUFA. Srebps are known to work with other factors such as nuclear transcription factor Y (NF-Y) and Sp1 in regulation of cholesterol and fatty acid biosynthesis in vertebrates (Reed et al., 2008). Within the snakehead *elovl5* promoter −161/+46bp, the SRE-like element is in close proximity with four putative Sp1 binding elements. These G-rich sequences are ubiquitous regulatory elements in many genes and have been shown to cooperate with Srebp in regulation of lipid metabolism (Reed et al., 2008). Sp1 binding sites was found in *fads2* promoter of several teleost species with significant Fads2 activities, such as zebrafish, salmon and rabbit fish but missing in several marine carnivorous counterparts, leading to the speculation of Sp1 having an integral role for *fads2* expression (Zheng et al., 2009, Geay et al., 2012, Xie et al., 2018). We now showed the presence of several putative Sp-1 binding sites in the promoter of a freshwater carnivorous teleost *elovl5* with validation of their requirements for basal transcription. Mutating one of the Sp-1 sequence in the snakehead −161/46bp promoter fragment lowered the transcriptional activities of the fragment. Intriguingly, mutation of one of the G-rich binding site significantly increased the transcriptional activity of the snakehead *elovl5* promoter, giving hint to a repressor function situation within this region. Besides exerting additive or synergistic effects on gene activation, other Sp1-like transcription factors such as Sp3 could also repress transcription (Bouwman and Philipsen, 2002). Taken together, these results insinuate a role for Sp-1 binding in the control of Fads or Elovl in species with significant LC-PUFA biosynthesis capacity.

Both snakehead and zebrafish *elovl5* promoter fragments with transcriptional activities in luciferase assay showed transient reporter-driven expression in YSL of embryos. Besides storing most of the lipids prior to transportation to support developmental processes, the teleost yolk is also an active site for lipid metabolism (Fraher et al., 2016, Pirro et al., 2016). The YSL hydrolyzes complex lipids into fatty acids and distributes them to the developing embryos with lipoproteins, before eventually degrading upon the exhaustion of yolk storage (Carvalho and Heisenberg, 2010, Kondakova and Efremov, 2014). Elsewhere, disruption of YSL-localized genes encoding for lipid metabolism and transportation enzymes resulted in perturbed yolk lipid utilization and transportation (Schlegel and Stainier, 2006, Chang et al., 2016). The presence of *elovl5* promoter signalling in YSL recapitulates the expression of *elovl* in YSL at 24hpf (Tan et al., 2010). Furthermore, the *fads* promoter was also localised in zebrafish YSL (Tay et al., 2018). We speculate that both these enzymes are required in the YSL to facilitate *de novo* synthesis of lipids, where fatty acids are repackaged into preferred lipid classes for distribution (Pirro et al., 2016). Collectively, these results suggest a Srebp-mediated role for *elovl5* in teleost yolk lipid metabolism and transportation.

In conclusion, we successfully isolated a 2497bp upstream of the snakehead *elovl5* promoter and −2592 bp upstream of the zebrafish *elovl5* promoter. Promoter fragments with basal transcriptional activities from both species require SRE site for transcriptional activation and were able to direct the expression of *elovl5* in zebrafish YSL, at stages where lipids are actively transported from yolk. In addition, a putative Sp1 binding site was also shown to have transcriptional activation role in the snakehead *elovl5*.

## Acknowledgments

This work was supported by the Universiti Sains Malaysia Research University-Individual Grant Scheme. [304/PBIOLOGI/650869/K105].

