## Supplement 1-4 for "Sterol regulatory element binding protein and Sp1 is required for transcriptional activation of a freshwater carnivorous fish (*Channa striata*) *elovl5* elongase"

**Supplementary 1.** Primers used for promoter studies, mutagenesis and EMSA experiments. The restriction sites are underlined, and the mutated sites are in bold.

|  | **Snakehead** | |
| --- | --- | --- |
| **Application** | **Primer** | **Sequence (5’ to 3’)** |
| **Promoter Cloning** | | |
| 1st fragment | CsElo5 gw R1  CsElo5 gw R2 | GGACCAATCCATGACTCAAAGAAAGTG  CAGTTTATGATTGATAGTCTCCATTTGTC |
| 2nd fragment | CsElo5 gw R3  CsElo5 gw R4 | AGGTACACGCTGGACTCATCCATTCTA  AAGCAAACAACACACCGAAGCATAATC |
| 3rd fragment | CsElo5 gw R5  CsElo5 gw R6 | AGTGATGACAGATGGCTGTGCCTAAAT  TAAGTCTCAGGGCTTTGTGTGTGCTTT |
| **pGL3 luciferase promoter-reporter plasmid CsElovl5** | | |
| -2497/+46 | CsElo5-2497  CsElo5+46 | GAGCTCTCTCTGGAAATCAAGTCTGTGTG  AGATCTCGATCGGCAGTGAGTAACCT |
| -1460/+46 | CsElo5-1460  CsElo5+46 | GAGCTCGAACCTTTCCCACCACAAGA  AGATCTCGATCGGCAGTGAGTAACCT |
| -1067/+46 | CsElo5-1067  CsElo5+46 | GAGCTCTTTGTTCTGGTCCCAAGAGG  AGATCTCGATCGGCAGTGAGTAACCT |
| -474/+46 | CsElo5-474  CsElo5+46 | GAGCTCAGTTTATGCTGCCCAACA  AGATCTCGATCGGCAGTGAGTAACCT |
| -295/+46 | CsElo5-295  CsElo5+46 | GAGCTCTGGCTCCTACATGAGTAAAGTGG  AGATCTCGATCGGCAGTGAGTAACCT |
| -161/+46 | CsElo5-161  CsElo5+46 | GAGCTCTCACAGAAGCCATCCAAAAA  AGATCTCGATCGGCAGTGAGTAACCT |
| **Site-directed mutagenesis pGL3-CsElovl5 -161/+46** | | |
| Mutate SRE | CsElo5 mSRE F  CsElo5 mSRE R | CAAGGCGGCACATTGTGCT**TTTTT**GGGGGCCGGACAGCCAAG  CTTGGCTGTCCGGCCCCC**AAAAA**AGCACAATGTGCCGCCTTG |
| Mutate Sp-1a | CsElo5 mSp1a F  CsElo5 mSp1a R | CTATTTTCCACCGAGCGACAA**TTTT**GCACATTGTGCTCAGACGGG  CCCGTCTGAGCACAATGTGC**AAAA**TTGTCGCTCGGTGGAAAATAG |
| Mutate Sp-1b | CsElo5 mSp1b F  CsElo5 mSp1b R | GCACATTGTGCTCAGACGGG**AAAAA**GACAGCCAAGGTTACTCAC  GTGAGTAACCTTGGCTGTC**TTTTT**CCCGTCTGAGCACAATGTGC |
| Mutate Sp-1c | CsElo5 mSp1c F  CsElo5 mSp1c R | CCGGACAGCCAAGGTTACTCA**AAAAA**GATCGAGATCTGCGATCTAAG  CTTAGATCGCAGATCTCGATC**TTTTT**TGAGTAACCTTGGCTGTCCGG |
| **GFP reporter plasmid construction** pZsGreen1-1-CsElovl5 | | |
| -2497/+46 | CsElo5 GFP -2497  CsElo5 GFP +46 | CCAACTCGAGTCTCTGGAAATCAAGTCTGTGTG  TTAAGGATCCCGATCGGCAGTGAGTAACCT |
| -1067/+46 | CsElo5 GFP -1067  CsElo5 GFP +46 | CCAACTCGAGTTTGTTCTGGTCCCAAGAGG  TTAAGGATCCCGATCGGCAGTGAGTAACCT |
| -474/+46 | CsElo5 GFP -474  CsElo5 GFP +46 | CCAACTCGAGAGTTTATGCTGCCCAACA  TTAAGGATCCCGATCGGCAGTGAGTAACCT |
| -295/+46 | CsElo5 GFP -295  CsElo5 GFP +46 | CCAACTCGAGTGGCTCCTACATGAGTAAAGTGG  TTAAGGATCCCGATCGGCAGTGAGTAACCT |
| -161/+46 | CsElo5 GFP -161  CsElo5 GFP +46 | CCAACTCGAGTCACAGAAGCCATCCAAAAA  TTAAGGATCCCGATCGGCAGTGAGTAACCT |
| **Zebrafish** | |  |
| **Application** | **Primer** | **Sequence (5’ to 3’)** |
| **Promoter Cloning** | | |
|  | Elo5-RACE-Primary-Rev | TTATACGGCACCAGCAGGGCTCTGCAAGAG |
|  | Elo5-RACE-Nested-Rev | GGGAATGTAGTCGTCCAGCAGAAACCATCC |
| **pGL3 luciferase promoter-reporter plasmid ZfElovl5** | | |
| -2592/+202 | Elo5-2K-Kpn1F  Elo5-2K-Xho1R | TTGGTACCCTCAAGCCGGAGTTTACAGC  TTCTCGAGGTATCATCCGCATTCGAGGT |
| -1286/+202 | Elo5-1.5K-Kpn1F  Elo5-2K-Xho1R | TTGGTACCTTAATCTGGGGTTGCCACAT  TTCTCGAGGTATCATCCGCATTCGAGGT |
| -682/+202 | Elo5-1.0K-Kpn1F  Elo5-2K-Xho1R | TTGGTACCATCGCCTCGTCGTAAAGAAA  TTCTCGAGGTATCATCCGCATTCGAGGT |
| -292/+202 | Elo5-0.5K-Kpn1F  Elo5-2K-Xho1R | TTGGTACCCACTTGCGTTCTGTTGAGGA  TTCTCGAGGTATCATCCGCATTCGAGGT |
| **Site-directed mutagenesis pGL3-CsElovl5 -161/+46** | | |
| Mutate SRE-P | Pro5-MutF  Pro5-MutR | ATCCTCGGATGTGTGTGGAAA**CCCC**AGCACGCGGGTGACCTCGAA  TTCGAGGTCACCCGCGTGCT**GGGG**TTTCCACACACATCCGAGGAT |
| Mutate SRE-D | Dis5-MutF  Dis5-MutR | TGTAACTTCAGATGAATGGG**CCCC**TTTACTTGAAGAAACAGTTC  GAACTGTTTCTTCAAGTAAA**GGGG**CCCATTCATCTGAAGTTACA |
| **GFP reporter plasmid construction** pZsGreen | | |
| -682/+202 | GFP-Foward-Xho1  GFP-Reverse-BamH1 | AT**CTCGAG**CTCGTCGTAAAGAAACAGTC  TT**GGATCC**GTATCATCCGCATTCGAGGT |

**Supplementary 2. DNA sequence of snakehead fish (*Channa striata*) *elovl5* promoter) and the schematic presentation of the promoter showing location of the three putative transcription start site.**

```
-2497	 ATCAGATGTG TATTTATTTT TCTCTGGAAA TCAAGTCTGT GTGAGAATGT AAATGTAGCA	 	-2438
-2437	 GAGAGTTGTA CAGTAGTCTG CTTTTAATCA TGTGTTCTGT AAATACTGTA CCTCTGCTCT	 	-2378
-2377	 ACATTAACAA GTTTTGTTTT TCTTGTTGAA CATGAGCTTT GTTTGCATTT GACAGAATTA	 	-2318
   -2317	 TTTTTGCTGC ATTTTAAGAG AAATGTATGA TCTGCTGACA TTGTAGTGGG ACCTTGAAGG	 	-2258
   -2257	 TAGTGTTTCA TTCAAATGGA AATGTCGAAC CATTGTTTGC AATGTGATGC TGCTTCAAAA	 	-2198
   -2197	 TAAATCTGTA AAACTGTAAA TTAAGCAGCT TTCATAGGTT TTATGGAACT TAAATACACC	 	-2138
   -2137	 CTGTGTCTGG TCTCTGTTAA TTGGGTGTCT TATTAAAGAA CATAAGAGCC ATGCAGGGCT	 	-2078
   -2077	 GGTAGGGTTG ACATTATTTC TTCAGGCATT TGTTTGAAAA TTAAAAGCCT TGGACAAATT	 	-2018
   -2017	 TACATTTATG CCTGATGATA TAATTGGTCC TTATGAGGCC CTAAAAGCTT ATTGCAATTT 	 	-1958
   -1957	 ATCAGTTGCA CCAGGTGGAG CGACGGCCTG ACCTTACTGT CCCCGGACAC AGGGGGGTTT	 	-1898
   -1897	 GTGAGGGAGG AAAGTAGCTT CAAAGAAGAT AAAGTTGGAG ATGTAATTAC AATTCTCTCC	 	-1838
   -1837	 ACTACCCACT TTGACTTCAA AGATCAACGG CTTTTGGCTT GTTCCTTCAT GGGGCACCAC	 	-1778
   -1777	 AGCAGAACAT GTTCCACACG CTGATTTGGC ATGAGATGCC CGTCCTGCCG CAGCCCTCCC	 	-1718
   -1717	 ATTTTATCCA GGCTCAGGAT CGGCACCTTG GTGGCTGGGT GGGCCACACC CGGTGGGGCG	 	-1658
   -1657	 GGAATCAGTC CCGCAGCCTT CTGCTTCCAA ACCCGACGCT CTACCCATTG AGTGACCTCA	 	-1598
   -1597	 ATAAGAAACA CATAGTAGAA TTATCATCTT TCAAACAGTT TTAACCTTAA AACTGAGAAA	 	-1538
   -1537	 TGATGGATTG CATTAAATTC CACAGGTGGT CATAATGTTA TGCCCGATTG GTGTATTTAC	 	-1478
   -1477	 CTACACTTCT TTTGTAACAC TTGCAAAAGA CAAACCAGAA CCTTTCCCAC CACAAGATGG	 	-1418
   -1417	 TGGTATTAAA CTGCAGCCTT ACATAGAAAT AAAAAGCCAC AGTTTGACCT CTGGGACCAG	 	-1358
   -1357 	 ATAAAGATTA TGGGCCTACC CTGTGAGGGA GGGATGAGAC ATACAGTAGT TTTAAACTGC	 	-1298
   -1297	 AAAAGAAATG AAAGCACACA CAAAGCCCTG AGACTTACAG TTGTTAATAC TGTGAGACCG	 	-1238
   -1237	 TAGTAATAAG AAAGCACAAA ACAATATAGT TTAATGTTAA GTACATTATG AGCGTGTTAG	 	-1178
   -1177	 ATAAGAACCT GAGGGAAAAA AAGTAAATAT GTTGCAGGTC AGTGAATTTA GGCACAGCCA	 	-1118
   -1117	 TCTGTCATCA CTGATCCCTG GAGACTTGTC ATAAAAATCA CACGGGTTAA CTACCAACAT	 	-1058
   -1057	 TAATAATGTC TTTGTTCTGG TCCCAAGAGG CGATCACAGA GGGATCAAAC TGAAACTCAG	 	-998
    -997	 	ACTGATTAAA TCCTGCCACT AGTTATGTTT TGTAAGTCCT CCTGTGCTTT TCTTATACTA	 	-938
    -937	 	AAAGTCAGTG ATCAGAGCAG ATCAGACTTA GGTTCGGGTG TGTCACAGGT AATGTGCAAT	 	-878
    -877	 	GTGGTCTCCA GTCAGCTTGT GTGAGCATTC CTATACTTCT GGCTCCTACT CTCACCCTGC	 	-818
    -817	 	CAAAAGTACA GTAGATACCC AGCAACCTGA TATCTAGACT GTATCTCTCT TGGGCTAAAG	 	-758
    -757	 	AGCCATGACC GTCCCGACTG GCTTGATAAG CAGTATCAAA TGAAACCAAC ATCTGCTCAG	 	-698
    -697	 	CAGTTCAGTC AGTCCTCAAC CCAACTTATC TGGCTCCACG TGTTACAAAT AGCTACTTGT	 	-638
    -637		 TCTATCAGAC GTCACCATGG TCCCACCTTT ACAGCCTCCC TGAATCCTCA CCCAGGCAAA	 	-578
    -577		 GAGCAGGTAC CCGGCTGAGC AGGTTCAACT ACAGCGCGTT GGTACCGCAG TCTGACCACA	 	-518
    -517		 ATCACCAGCT GGTTTCAGTA GTTTGTGGAG TTAAACTGCC CTGATTCACA GAGAACTGAA	 	-458
    -457		 TCCAGTTTAT GCTGCCCAAC ACCAAATGAC AAATTGGACT AATGACTCCA TTCACTGAGA	 	-398
    -397		 AAGTGACACA AGTGACTGAC CAGGTTCAGT AATGAAGTCA ATAAAATTAG AGGAGAAAAC	 	-338
    -337		 GTTATTGCAG ACTTTATGCA TATATACTTA CGATGCATTG CATTGTTTCT AAATGATTAA	 	-278
    -277		 ACTGGCTCCT ACATGAGTAA AGTGGCATTT ACATGTTTCT TCCCAGTGGT TCCATTGTTC 	 	-218
    -217		 TTAAATCCAC GAGCACAGTC ACTGACCCTA TGTGCTCCCT GCACATACAT TCATGTGCCT	 	-158
    -157		 CCGGTTAAAA ATTAATTCAC AGAAGCCATC CAAAAAAAAA ACCTTTGATG TCATCTTTTA	 	-98
     -97		  TGACATCGTT GCAACCCACC GGGTCACCTA CTGGCTGCAG TCCTCCTGCA CGGCAATGGG 		-38
     -37		  GCCAGATTGC GTGTTCGTCC TCCCACTATT TTCCACCGAG CGACAAGGCG GCACATTGTG	 	+23
     +24		  CTCAGACGGG GGCCGGACAG CCAAGGTTAC TCACTGCCGA TCGCTCGCCC CGCCTCGAAA		+83
     +84		  GGTAAGACGC TGCCGGGGCT GCATCTCCTC ATCCACACAC ACACACACAC ACTCACTCAG    		+143
    +204		  ACATTTTCAG GGCGCGCATG GCGTCTGGGT GATGCTGAGG CGACGAGCTC CCGAGCTACC	 	+263
    +264		  GCGCGATCGC CTGGACAAGT TCGTGCGTTG TTGTTATTAT TATTATTATT ATTATTATTA		+323
    +324		  TTAATCATCA TTCGTTTCTG CCTCTTTCGG GAATTGCACG GCCATGTAGT CGACGTCAGC	 	+383
    +384		  TCCCGTCTTT GAGATCCACC GGAGATGATT TGATCACACG TGTGACCGTG ATGGAAAATC		+443
    +444		  ACTATTATAC AATAGTGAGG CGAAGAGAAG CGGCGTGTGG CGCGATATTC CCGGCGGATT	 	+503
    +504		  TCTCCTGCAT CCCGCCGATC ACAACATCCT ACAATATTCC CGTGAAGACA CCGAGCAGGA	 	+563
    +564		  ATATTGTACC TACTGTAAAT GATTTCTAGT TGTCAAACTT GTTAGTGTTG GTCCTGCTGA		 +623
    +624		  AAACTTTTCG TTGCTGCTAT TGCTGCTGTG ACTGACTGTG ACTTTGCACT TTCACATGCT		 +683
    +684		  TTAATTTGTG CACACGTTTT TAGGCCAGCA CAGACATAAA TCTAATATTA ATTATTCTGC		 +743
    +744		  CTGTCCGTGG ATGTGATTGT CTGTGCACCA GTTAGCCTCA TCTACAGGGT CATGCGAATG		 +803
    +804		  CAGGACAATG TGCAGCCTAA ACTGGACTTT GATTTATCAT TTTATGTATC AGGGTTGGCG		 +863
    +864		  GATAAACCCA GCTTTCACTT TAGTGCAGTT CTGTCTCTCT AACCCTCTGT CATGTGTTTG		 +923
    +924		  CAATTTAAGA CCATGTCAAT GCAGATTAAA CTACACAATT AATAGAAATG TAATAACAAC		 +983
    +984	  	ATTCCTGCAC ACCGGTATGA CAAGTATGTC TTCATAAACA AATTTATCCG TCCATAAAAT		 +1043
   +1044	  ACCAACTCAC TCTTGAGTGA TTTGATTTGC AGCATTACCC ACAATGGCCT GATGCATCCA		 +1103
   +1104	  GGTTTCAGAC GGCCACATAC CACATCCAGG ATTGCAAAAT CTCTGAGAAC ACACTCATTT		 +1163
   +1164	  CACAGTGATT CATGTTATTC ACTATATCAT CATTATGCCC ATCCCGAAAC GTTACAGTAA		 +1223
   +1224	  ATAAGACAGC TATAAGAAAT CATTATCACA TGTGGTGTAT TGCTGTCCAA AGTCCAGATT		+1283
   +1284	  ATGCTTCGGT GTGTTGTTTG CTTGTTTAGA ATGGATGAGT CCAGCGTGTA CCTCGCAGAG		 +1343
   +1344	  ACCCACTGAC TTCCCCCTGT GGTGGCTTAT CATGCTCTGA TTAAGTCCGT ACAGATTATT		 +1403
   +1404	  ACGTTTTGGC GCAGCCTAAT GCATCCTGAT AATCAGAGCA TGGATATTAG TCGATTGGGC 		 +1463
   +1464	  TTGATCAGAA TAATCCATAT TTATTTCCAG CCGCTAAGAC AACCGAAGAG AAATGTGTTA		 +1523
   +1524	  TTGTTGCCTA GGTAGCAGTT CACTGCAACC TGCAGGTTTT AACACAGCCT GACCTGCAAA		 +1583
   +1584	  AATTCCTTTG TAAACTATAC ACATTGCACA ATATTTAAAG AGAGTTTTGA GGACTTTTGG		 +1643
   +1644	  GCTGTGCTGT GCTCCTCGCA CGCCGTTGTG CAACTGAAAT TAATTTGACC ATTTAAAGGA		 +1703
   +1704	  TTTACGTTAA TATTTCTTTA TATGGATTTT AGAATATGAA GTTGTGTAAA CAAAGTCCTT		 +1763
   +1764	  GCGTTGACCC CGCAGTTCTA ATAGTCATAT TTCAGTATCA GTTTACGGTT TAAACGAGCC		 +1823
   +1824	  CTGTGCCGGC TGAATTGAGA TGCTTTAGCT GAATTTGATA GATCTGCTCC AGTTTCTGTC		 +1883
   +1884	  TGTAAAGTGC AGGGCTGATG TGTTCACCTG TAACGCTGTG CTGTACTTTC AAATCAGACA		 +1943
   +1944	  CACACTCCTG CTTCGTTAAC TTCAATTTCA GTTCAGGACA TTGTGTCTTT ATTTAAGAGA		 +2003
   +2004	  CCAATGTGGC CCTCCATTAT TTATGGAAAC CATAATATAC CAGGTCTGAA GGTGGTATTA		 +2063
   +2064	  TTGAAGAAAG ACTCACACCG TCCTCCGAGT CTTTAGTTTG TCACGTGTTA TATTTCTAGC		 +2123
   +2124	  AGTAGGCGCA GGTGTAGAGA CCGTCCAACG GTGCAGCAGT GTGACGTAAC TGGACTGAAT		 +2183
   +2184	  TGTATGTGTA TCTCAGACAC ATGTCACTGC AAATAAAGAA CGCAGACATT CAATGATTTC		 +2243
   +2244	  AGCGGCTGGG GGGGAGGCAT TTTGTGTAAA GGCTTAAGCT CTATATTATG GGATGTTTGT		 +2303
   +2304  	  TGTAAGAAAG ACTTTGTGTG AGACAGATTC ACTGTTCACT GCTGTTACAA CACTGAAGGG		 +2363
   +2364	  ATTTTTTAAA AGCCTCCTCT GCTGTAAAAA CATGACTGTG CAACTTAACG TGTTTGTCCA		 +2423
   +2424	  GGCTACGTTT TCAGTAAAAA TACTTCCTGT AGTTTTAGGA TATGTTAGTA GGAGAGTTGA		 +2483
   +2484	  CAAGCATCTC TGAGAACTGA ACCCAGTAAC TTTAGATATA TATTTATAAG GAAAATTATC		 +2543
   +2544	  CAGCTGTGAA CATCCAGACA CTCTCACAAT CTGCCTTACA CCACAGGGTT TATCTGACTA		 +2603
    +2604	 TCAGGGCAGT GGAGTCTAAC CTTCTATTTT AACTTGCGCA CACAGTTTAT CAGAAAGAAG		 +2663
    +2664	 TACTTAGACT CTCCGTGGGA AAATCCAGAG AAAAACATCT ACTTTTCAGT CATCCCCGAC		 +2723
    +2724	 AGTTGCACAG GTAGCATGTG TTTGAATCGC TCACTAAGAC AGAGCGGTCA GCTGAGAGAA		 +2783
    +2784	 TCACTAACTC ACCACGATTT GGAGCCCACG TAAGAAAACG TGGACAACAA ATGGAGCAGT		 +2843
    +2844	 GCTGCCTGCC TTGTTTGAAA CCACAGGAAA GATTGATAAT TAAACACATA ATACTATAGA		 +2903
    +2904	 TAGAGAGGAT GACACAAGAC TAATGAGCAA GTAGCATCAT GACTATGAAA AAGTTATGAT		 +2963
    +2964	 TGCAGCTGCT GTCTGTAATA GTAATCACTG GCATCAATCA ACCTCCTTTT TTGAATCATA		 +3023
    +3024	 ATTGAGCTTA TCATAGTGAA GCTGCACTAA TCACCATTCT TACGAAAAAC AAACTCGTAG		 +3083
    +3084	 CGTACCACCC TTTTTCCGAG TTCCCCTCAG AACATTTCAC CATTTTTGAC TTTACTGTGA		 +3143
    +3144	 CTCACTCACT CATGCTGCTC CCATCAGTGT TACCACCTGT TGCAACAAGC TCCGAGAACA		 +3203
    +3204	 CTGTACACAA CCGGCTCAGC ACCAGACGCA GACTGGCAAA GTTATCAATG ATGTGCACAT 		 +3263
    +3264	 TTATCGGTTA CGGATACACA CATTTCTCTC AGCAGTTGGT GGAGGCAAAG GCAGAGCTAA		 +3323
    +3324	 CAAAGAGAGT TTCACTTTAA AAGTTGTAAA TAATTGCCTG TATTTTGTGT TTAGGTGACA	 	 +3383
    +3384	 AATGGAGACT ATCAATCATA AACTG                                      	 									 +3408
```

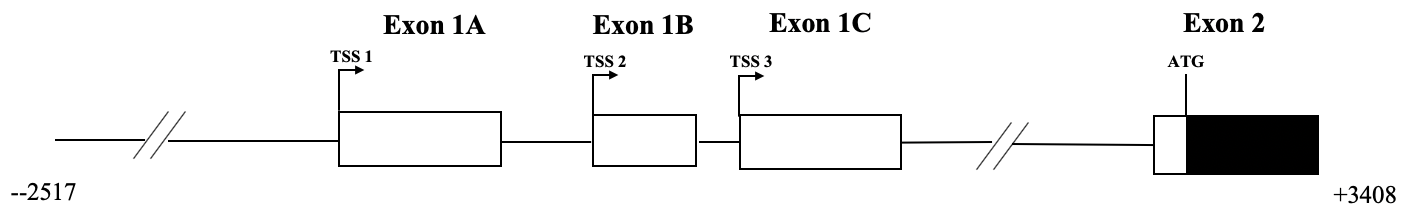

**Supplementary 3. DNA sequence of zebrafish (*Danio rerio*) *elovl5* promoter and the schematic presentation of the promoter showing location of the two putative transcription start site.**

```
-2592 	GTGTTGCTTG TTTGCTTCAG TCTGCTTGCT TAACCCATCA GGTGTTTTAA GTCAGCAGTT		-2533
-2532	TGATTTTCTT TCATTTTTAG AGATAAAATG TAGGTCTTTA TGTTTTAAAT ACAGCTTGGT		-2473
-2472	GGGTGTTTGC TAGTTCTTCA TGTTTATTAC AGTCAAACGG AAATCCATCT TTAAAGTACA 	-2413
-2412	ATAGAAGTGA GTTATATATT AATAAAAGTA AACTATAAAT ATATATGTTA ATAGAAGTGA		-2353
-2352	GTTATATTAA TGTAGATAGA TAGATAAATA GATGGATAGA CAGACAGACA GATAGATAGA		-2293
-2292	TAGATAGATA GATAGATAGA TAGATAGATA GATAGATAGA TAGATAGATA GATAGATAGA		-2233
-2232	TAGATAGATA GATAGATAGA TAGATAGATA GATAGATAGA TAGATAGGCA GACAGACAGA		-2173
-2172	CAGATATGTG GATGGATGGA TAGATAGATA GACAGACAGA CAGACAGATA TATGGATGGA		-2113
-2112	CGGATAGATA GACAGATAGA TAGATAGATA GATAGATAGA TAGATAGATG GATAGATGGA		-2053
-2052	TAGATAGATG GATAGATAGA TGGATGGATA GGTAGACAGA CAGACAGACA GACAGACAGA		-1993
-1992	TATGTGGATG GATGGATGGA TGGATGGACG GACGGACGGA TAGATAGATA GATAGATAGA		-1933
-1932	TAGATAGATA GATAGATAGA TAGATAGATA GATAGATAGA TAGATAGACA GACAGATATG 	-1873
-1872	TGGATGGATG GATAGATAGA CAGACAGACA GACAGACAGA TAGATAGATA GATAGATAGA		-1813
-1812	TAGATAGATA GATAGATAGA CAGACAGATA GACAGACAGA CAGATATGTG GATGGATGGA		-1753
-1752	TGGATAGATA GATAGACAGA CAGACAGATG GATGGATGAT AGATAGACAG ACAGACTGAT		-1693
-1692	GATGATGATG ATGATGATGA TGTAGATTGA TGATGGATGA GACGCGATGA CCACGATGTA		-1633
-1632	TCTATGACGT ACAGTCGCTA GCCTATCGTC AATGATGCGC ATTCCATGTG GCTGACTAGT		-1573
-1572	GCAGTCTGTA CATCTATCCA TCATGCATAT GACTGTCGAT TGATCGCTAG TGAGTAGACT		-1513
-1512	GTAGTAGTAG ACGTAGTGTG CTATGTGCAC GCAGCGATTG ATGATGATGG TAGTAGCAGA 	-1453
-1452	CGACCGACCA GATATTGATG ACGATGGATG ACGATGATGA TAGATGATGA TAGATAGATG 	-1393
-1392	ATGATGATTG ATAGATAGAT GATAGATAGA TGATGATAGT AGACAGACAG ACAGACAGAT 	-1333
-1332	AGACAGATTT CGGCTTAGTC CCTTTATTAA TCTGGGGTTG CCACATCGGA ATGCCCCGCC 	-1273
-1272	AACTTATCTA ACATATGTTT TACACATTGG AACCCTTTCC AGCTGCAACC CAGTACTGGG 	-1213
-1212	GAATACCCAT ACACACTTAT TCACACACAT ACACTACGGC CAATTTATTT GACCCGATTC 	-1153
-1152	CCCTATAGTG CATGTGTTTG GACAGTGGGG GAAACTGGAG CACCCGGAGG AAACCAACGC 	-1093
-1092	GGGGAGAGCA TGCAAACTCC ACACAGAGAT TCCAACTGAC CAACCTGGGC TCGAACCAGC 	-1033
-1032	GACCTTCTTA CTGTGAGGTG GCAATGCACT GAGCCACCAT GTCGCCCCAT TCATTGACAA 	-973
-972		TGTTTTGTAA CTAATATATT AAATAGGCCT AAAGTATAAT GCACTATATA TTACAGTAGG 	-913
-912		GTTTAAAAAC CGCAACATTT ATTACAAGTT TATACAAGTT CGTAGATTGT TTTTTAAGAA 	-853
-852		GTAAGCCATC ACCTCCAAAT CAGATACGCC GTCTGGGTCA AAGTCTTTGT ATGTTTTAAA 	-793
-792		TTGTGTGTAT GAAATAAAGG TTTATAAATG TGACGCTGAA GTGTGTGTAG GGGTTTAGCA 	-733
-732		CATGATGCCA CAAAGTAATG GGACTTTGTA TCGCCTCGTC GTAAAGAAAC AGTCTGTTCT 	-673
-672		CTGACCAGAT TACCTCCAGA TAAATTTGAT TCAACTTTAT GACCTCATTC ACCAAAATAC 	-613
-612		CGCATACACA TTGACAAGAA CTCGGCATTG AAAGCACAGC AGCTCTATAA AGGCAGTAAT 	-553
-552		TTTTAACCGA ACTCTCTAAC TCCTGCGATA GTCTGATTCC CTTTATCACC TGCTGTCATG 	-493
-492		CGCTCTTTGA TTGGACGGAT CTGGTTTCAC ATGAGAGCAG CTCAGCAATT GGTCCTTTTT 	-433
-432		AGTTTCACAT GAGAGCAGTG CTCTGATTGG TCTGTTGTAA CTTCAGATGA ATGGGGTGAT 	-373
-372		TTACTTGAAG AAACAGTTCA CGCTTTGTAA AATGTGAACG TGCAGAACAG CTGCCGGAGC 	-313
-312		GCGCGCTTTT ATCTTAGAAC GTCACATTCT AAAACTACAC ACTTGCGTTC TGTTGAGGGG		-253
-252		GAATTGATAC ATTCGCGTTC TAAATCAACT TCAAACATGA GTACAGTTGC ATGACGTCAT 	-193
-192		CATAAATGTC AACAAACTGC TACATCATTA AAAGTTTGTT GCATCTCCTC AGCGCCTCCT 	-133
-132		GGTAAAAATC GAATGAGGTG TGTGTGAAGT GTTTAACGTG TCTCGGATCG ATGTGTGTGT 	-73
-72		GTGTGTGTGT GTGTGTGTGT GTGCTCGCAG ACTTTGCCCT GAGAGTCTTC AAACACGTCC		-13
-12		CGTTATGCGC ATCAGATGAA GGTGGACTAC ACACACTCAT GACGCTCATT GGTTCAGCAG 	+49
+48		CTTCACACTC AAACACACAC ATCCTCCAAG GACAGGACGA AGCCCTGAGC AACACACACT 	+107
+108		CGCACACTTC TAAAGACTAA AGGTAAGATG CAGATGAATA TTCTCGAATT TTGTAAACAT 	+167
+168		CCTCGGATGT GTGTGGAAAG TGAAGCACGC GGGTG  							+202
```

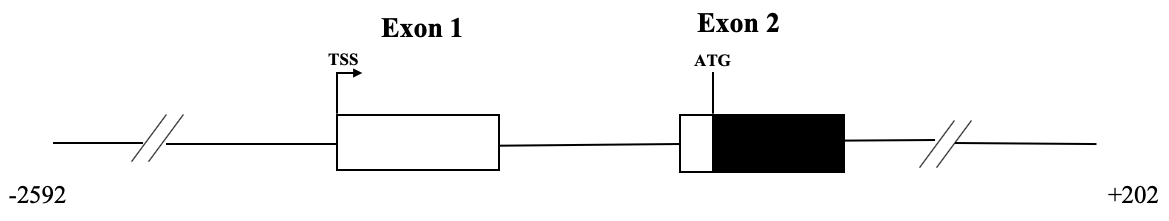

**Supplementary 4**. Nucleotide sequence and putative transcription factor binding sites of the (A) snakehead *elovl5* promoter (-161/+46 bp). The putative binding sites for Sp-1 and Srebp transcription factors are bolded and underlined, respectively. (B) zebrafish elovl5 (-682/+202 bp). The putative binding sites for Srebp transcription factors are underlined. TSS are indicated with bended arrow.

**A**

-161 TCACAGAAGCCATCCAAAAAAAAAACCTTTGATGTCATCTTTTATGACATCGTTGCAACC

-101 CACCGGGTCACCTACTGGCTGCAGCCCTCCTGCACGGCAATGGGGCCAGATTGCGTGTTC

**Sp-1**	**SRE**	**Sp-1**

-41	GTCCTCCCACTATTTTCCACCGAGCGAC**AAGGCGGCAC**ATTGTGCTCAGACG**GGGGCCGG**

**Sp-1**

+20	**AC**AGCCAAGGTTAC**TCACTGCCGA**TCG

**B**

-682	AGTCTGTTCTCTGACCAGATTACCTCCAGATAAATTTGATTCAACTTTATGACCTCATTC

-622	ACCAAAATACCGCATACACATTGACAAGAACTCGGCATTGAAAGCACAGCAGCTCTAT

-564	AAAGGCAGTAATTTTTAACCGAACTCTCTAACTCCTGCGATAGTCTGATTCCCTTTATCA

-504	CCTGCTGTCATGCGCTCTTTGATTGGACGGATCTGGTTTCACATGAGAGCAGCTCAGCAA

-444	TTGGTCCTTTTTAGTTTCACATGAGAGCAGTGCTCTGATTGGTCTGTTGTAACTTCAGATG

**SRE-dis**

-383	AATGGGGTGATTTACTTGAAGAAACAGTTCACGCTTTGTAAAATGTGAACGTGCAGAAC

-324	AGCTGCCGGAGCGCGCGCTTTTATCTTAGAACGTCACATTCTAAAACTACACACTTGCGT

-264	TCTGTTGAGGGGGAATTGATACATTCGCGTTCTAAATCAACTTCAAACATGAGTACAGTT

-204	GCATGACGTCATCATAAATGTCAACAAACTGCTACATCATTAAAAGTTTGTTGCATCTCC

-144	TCAGCGCCTCCTGGTAAAAATCGAATGAGGTGTGTGTGAAGTGTTTAACGTGTCTCGGAT

-84	CGATGTGTGTGTGTGTGTGTGTGTGTGTGTGTGTGCTCGCAGACTTTGCCCTGAGAGTCTT

-23	CAAACACGTCCCGTTATGCGCATCAGATGAAGGTGGACTACACACACTCATGACGCTCAT

+37 TGGTTCAGCAGCTTCACACTCAAACACACACATCCTCCAAGGACAGGACGAAGCCCTGAG

+97 CAACACACACTCGCACACTTCTAAAGACTAAAGGTAAGATGCAGATGAATATTCTCGAAT

**SRE-pro**

+157 TTTGTAAACATCCTCGGATGTGTGTGGAAAGTGAAGCACGCGGGTG
